# Acrylate reductase of an anaerobic electron transport chain of the marine bacterium *Shewanella woodyi*

**DOI:** 10.1101/2024.02.11.579808

**Authors:** Yulia V. Bertsova, Marina V. Serebryakova, Vladimir A. Bogachev, Alexander A. Baykov, Alexander V. Bogachev

## Abstract

Many microorganisms are capable of anaerobic respiration in the absence of oxygen, by using different organic compounds as terminal acceptors in electron transport chain. We identify here an anaerobic respiratory chain protein responsible for acrylate reduction in the marine bacterium *Shewanella woodyi*. When the periplasmic proteins of *S. woodyi* were separated by ion exchange chromatography, acrylate reductase activity copurified with an ArdA protein (Swoo_0275). Heterologous expression of *S. woodyi ard*A gene (*swoo*_0275) in *Shewanella oneidensis* MR-1 cells did not result in the appearance in them of periplasmic acrylate reductase activity, but such activity was detected when the *ard*A gene was co-expressed with an *ard*B gene (*swoo*_0276). Together, these genes encode flavocytochrome *c* ArdAB, which is thus responsible for acrylate reduction in *S. woodyi* cells. ArdAB was highly specific for acrylate as substrate and reduced only methacrylate (at a 22-fold lower rate) among a series of other tested 2-enoates. In line with these findings, acrylate and methacrylate induced *ard*A gene expression in *S. woodyi* under anaerobic conditions, which was accompanied by the appearance of periplasmic acrylate reductase activity. ArdAB-linked acrylate reduction supports dimethylsulfoniopropionate-dependent anaerobic respiration in *S. woodyi* and, possibly, other marine bacteria.

## INTRODUCTION

In the absence of oxygen, many microorganisms are capable of anaerobic respiration using various organic or inorganic compounds as terminal acceptors in electron transport chain. The most typical organic terminal acceptor is fumarate, but the variability of catalytic amino acid residues in homologues of fumarate reductase in various bacteria suggests the existence of other 2-enoate reductase activities [1]. Enzymes with different substrate specificities, such as cytochrome *c*:methacrylate reductase, cytochrome *c*:urocanate reductase, NADH:(hydroxy)cinnamate reductase, and NADH:acrylate reductase [1-6], have been indeed identified among fumarate reductase homologues.

Expression of NADH:acrylate reductase gene in the marine bacterium *Vibrio harveyi* is induced by acrylate regardless of the oxygen concentration in the growth medium [1]. This finding suggests that the main function of NADH:acrylate reductase is acrylate detoxification in *V. harveyi* cytoplasm, rather than anaerobic respiration on this unsaturated carboxylic acid. On the other hand, the usability of acrylate as a terminal electron acceptor has been documented for the bacterium *Halodesulfovibrio aestuarii* (formerly known as *Desulfovibrio acrylicus*). As the enzyme(s) responsible for the acrylate reduction in the latter bacterium has(ve) not been identified [7], it is interesting to determine their relation to the known NADH:acrylate reductases.

The main natural source of free acrylic acid is dimethylsulfoniopropionate (DMSP) [8]. Many marine algae and plants accumulate this compound in cytoplasm in very concentrations (up to hundreds of mM) as a compatible solute. The overall production of DMSP in the Earth’s biosphere is estimated as approximately 10^9^ tons per year, making it a significant source of carbon, reduced sulfur, and energy for various marine bacteria. DMSP cleavage by bacterial DMSP lyases DddL, DddP, DddQ, DddW, and DddY yields acrylate (Fig. 1) [8], which explains the abundance of acrylic acid in some marine ecological niches.

**Figure 1.**
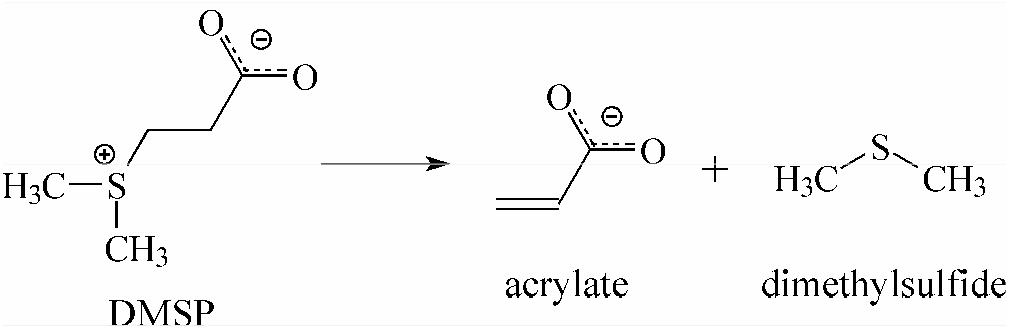
Dimethylsulfoniopropionate cleavage catalyzed by DMSP lyases.

DddY is specific among DMSP lyases in that it is highly active and has a periplasmic localization in the bacterial cell, which is unusual for DMSP lyases [9]. The periplasmic localization may indicate that the acrylate generated from DMSP is used in the periplasmic space rather than transported into cytoplasm, consistent with the presumed role of acrylate as a terminal electron acceptor for the anaerobic electron transport chain. Indeed, the genomes of many marine bacteria from the genera *Shewanella, Ferrimonas*, and *Arcobacter* ([8, 9], Fig. S1) contain, in a close proximity to *ddd*Y, genes encoding one or two copies of the flavin-containing and heme C-containing subunits of a hypothetical flavocytochrome *c* (Swoo_0275 and Swoo_0276 in Fig. 2a, respectively), similar to the cytochrome *c*:fumarate reductases of these bacteria. The products of these genes contain Tat- and Sec-type signal peptides for the flavin- and heme-containing subunits, respectively, indicating their periplasmic localization. The amino acid residues involved in the binding of the C4-carboxyl group of fumarate in fumarate reductases [1, 4] are conserved in *ddd*Y-associated flavocytochromes *c*. In contrast, the His and Thr(Ser) residues involved in fumarate C1-carboxyl group binding in fumarate reductases are replaced by Gly and Tyr in the *ddd*Y-associated flavocytochromes *c* (Fig. 2b). A likely corollary is that the DddY-associated flavocytochromes *c* are involved in the reduction of α,β-unsaturated carboxylic acid(s) different from fumarate.

**Figure 2.**
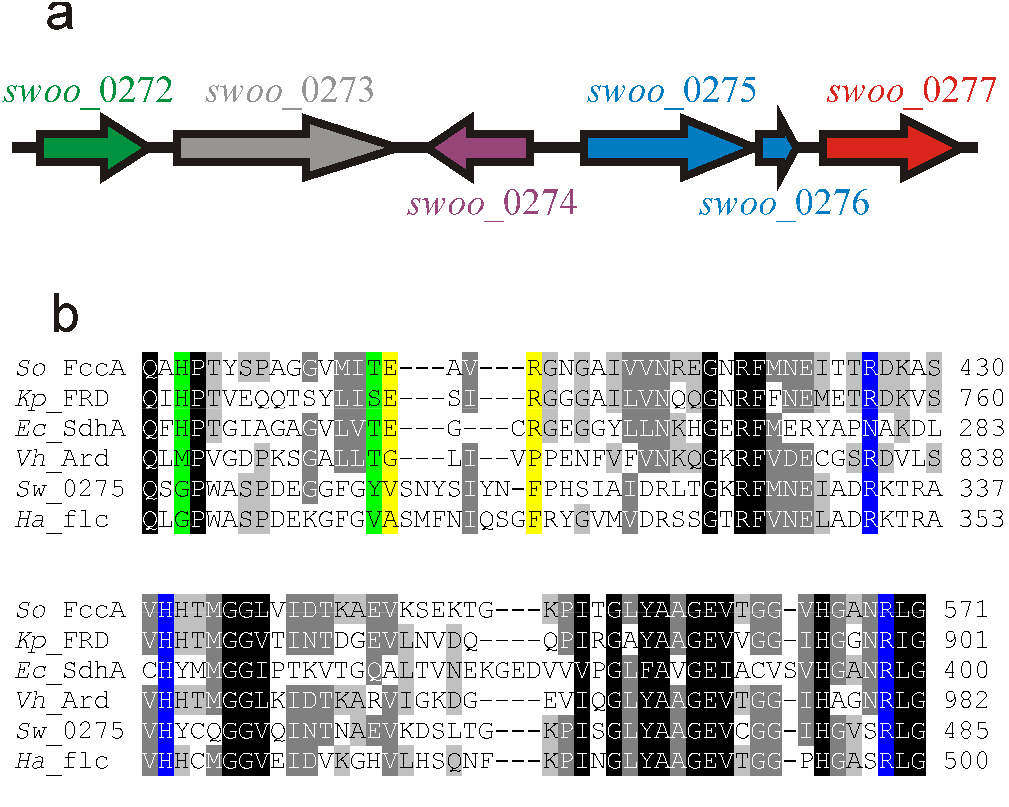
*ddd*Y-linked genes of marine bacteria. (a) The arrangement of the *ddd*Y-associated genes in the *S. woodyi* chromosome: *swoo*_0272, NADPH:acrylyl-CoA reductase; *swoo*_0273, hypothetical protein with unknown function; *swoo*_0274, transcriptional regulator; *swoo*_0275, flavoprotein subunit of flavocytochrome *c*; *swoo*_0276, tetraheme cytochrome *c*; *swoo*_0277, DMSP lyase DddY (https://www.kegg.jp/kegg-bin/show_organism?org=T00676). (b) Multiple alignment of the amino acid sequences of cytochrome *c*:fumarate reductase of *S. oneidensis* MR-1 (*So*_FccA, GenBank: AAN54044), NADH:fumarate reductase of *Klebsiella pneumoniae* (*Kp*_Frd, B5XRB0), an SdhA subunit of succinate dehydrogenase of *Escherichia coli* (*Ec*_SdhA, HDZ3930178), NADH:acrylate reductase from *V. harveyi* (*Vh*_Ard, P0DW92), a Swoo_0275 protein from *S. woodyi* (*Sw*_0275, ACA84576), and flavocytochrome *c* of *H. aestuarii* (*Ha*_flc, SHJ73509). The two fragments of the alignment shown contain the amino acid residues corresponding to the fumarate reductase residues involved in the binding of C4- and C1-fumarate carboxylates (blue and green, respectively) and in proton transfer to fumarate (yellow).

Many *ddd*Y-associated gene clusters (Fig. S1) contain in addition a gene encoding NADPH:acrylyl-CoA oxidoreductase AcuI (Swoo_0272 in Fig. 2a), which is responsible for the detoxification of cytoplasmic acrylate [10]. Acrylate metabolism may thus depend on the entire *swoo*_0272–*swoo*_0277 gene cluster, which makes the *S. woodyi* flavocytochrome *c* Swoo_0275/Swoo_0276 and homologous enzymes from other *ddd*Y-containing bacteria good candidates for acrylate reductases of the anaerobic respiratory chain. The aim of the present study was to test this hypothesis.

## MATERIALS AND METHODS

### Bacterial strains and growth conditions

*S. woodyi* cells were grown at 25°C in MR medium (31.5‰ sea salts (Marine Life, Russia), 20 mM L-lactate, 0.5% peptone, 0.25% yeast extract, 20 mM HEPES/NaOH (pH 7.5)) aerobically or anaerobically in the presence of appropriate electron acceptors. For Ard purification, cells were grown anaerobically in the MR medium supplemented with 20 mM dimethyl sulfoxide (DMSO) and 1.5 mM acrylate.

*Shewanella oneidensis* cells were grown at 28°C aerobically in LB medium or anaerobically in a medium containing 0.225 g/L K_2_HPO_4_, 0.225 g/L KH_2_PO_4_, 0.46 g/L NaCl, 0.225 g/L (NH_4_)_2_SO_4_, 0.117 g/L MgSO_4_·7H_2_O, 20 mM L-lactate, 0.05% yeast extract, 20 mM HEPES/NaOH (pH 7.2). When required, the growth media were supplemented with kanamycin (50 μg/mL).

### Construction of expression vectors

The expression vector for the ArdAB protein was constructed by amplifying the *ard*-operon (*swoo*_0275–*swoo*_0276) from the genomic *S. woodyi* DNA. A high-fidelity Tersus polymerase (Evrogen, Russia) and the primers Sh_wood_dir / Sh_wood_CR4_rev (primer sequences are listed in Table S1 in the Supplemented Information) were used. The resulting 2571-bp fragment was cloned into the pCR4-TOPO vector (Invitrogen, USA), yielding the pSwoo_0275&Swoo_0276 plasmid.

The expression vector for the ArdA protein was constructed by partial digestion of the pSwoo_0275&Swoo_0276 plasmid with *Hind*III and *Not*I restriction endonucleases. The product shortened by 350-bp was blunted and self-ligated, forming the pSwoo_0275 plasmid. The resulting plasmids were verified by sequencing and transformed into *S. oneidensis* MR-1 cells using electroporation.

### Preparation of periplasmic fraction from *Shewanella* cells

*S. woodyi* or *S. oneidensis* cells were harvested by centrifugation (10,000*g*, 10 min) and washed twice with a washing medium (0.5 M NaCl, 10 mM Tris-HCl (pH 8.0) for *S. woodyi* or 75 mM NaCl, 10 mM Tris-HCl (pH 8.0) for *S. oneidensis*). The periplasmic fraction was obtained by a Polymyxin B treatment [11, 12]. The cell pellet was suspended in the corresponding washing medium and supplemented with Polymyxin B (2,000 units/mL). This mixture was incubated in an ice bath for 20 min. The Polymyxin B-treated cells were removed by centrifugation (10,000*g*, 10 min) and the supernatant was used as periplasmic fraction for measurements of acrylate reductase activity and protein purification.

### Purification and characterization of Ard

Periplasmic fraction of *S. woodyi* cells was concentrated using a 30-kDa cut-off centrifugal filter, diluted with 10 mM Tris-HCl (pH 8.0) to 80 mM NaCl concentration, and applied onto a DEAE-Sepharose CL-6B column (16 × 30 mm) equilibrated with buffer 1 (10 mM Tris-HCl (pH 8.0)) containing 80 mM NaCl. The column was washed with three bed volumes of buffer 1 containing 100 mM NaCl, and Ard was eluted by a linear NaCl gradient from 100 to 340 mM in buffer 1. The protein that remained bound was eluted with 2 M NaCl in buffer 1. The most active Ard fractions were combined, concentrated on a centrifugal filter, and stored at –70°C. Protein concentration was determined by the bicinchoninic acid method [13] using bovine serum albumin as a standard.

Ard-bound flavins were extracted from the protein by trifluoroacetic acid and separated by HPLC as described elsewhere [12]. SDS-PAGE was performed using 12.5% (w/v) polyacrylamide gels [14]. The gels were stained for protein with PageBlue™ staining solution (Fermentas, Lithuania) or for heme C by 3,3′,5,5′-tetramethylbenzidine/H_2_O_2_ treatment [15]. MALDI-TOF MS analysis was performed on an UltrafleXtreme MALDI-TOF-TOF mass spectrometer (Bruker Daltonik, Germany) as described elsewhere [5].

### Determination of enzymatic activities

The acrylate reductase activity was determined at 606 nm by following the oxidation of reduced methyl viologen (MV, ε_606_ = 13.7 mM^-1^ cm^-1^) with a Hitachi-557 spectrophotometer. The assay was performed at 25°C in a 3.2 mL anaerobic cuvette. The standard assay mixture contained 100 mM HEPES/Tris (pH 7.5), 0.05–1 mM electron acceptor, and 1 mM MV. MV was pre-reduced with sodium dithionite until the absorbance at 606 nm of approximately 1.5 was obtained, which corresponds to the formation of ∼100 μM reduced MV. One unit of the enzyme activity was defined as the enzyme amount catalyzing oxidation of 2 μmol MV per 1 min.

The Michaelis-Menten parameters of acrylate and methacrylate reductase reactions were estimated from the integral kinetics of their complete reduction, as monitored by absorbance at 606 nm. Rates were estimated at 20 time points along the progress curve as the slopes of the tangents (-*d*[MV]/*d*t) using MATLAB (The MathWorks, Inc., USA). The residual electron acceptor concentration was calculated at each point from *A*_606_ using the MV:acceptor stoichiometry of 2:1 and assuming that the limiting value of *A*_606_ corresponds to 100% conversion of the electron acceptor. The Michaelis-Menten equation was fitted to the rate data using non-linear regression analysis.

The propionate dehydrogenase activity was measured at 600 nm in the presence of phenazine methosulphate and 2,6-dichlorophenolindophenol as electron acceptors. The assay mixture contained 100 mM HEPES/Tris (pH 7.5), 2 mM propionate, 2 mM phenazine methosulphate, and 25 μM 2,6-dichlorophenolindophenol.

### Induction of acrylate reductase activity and *ard*A and *ddd*Y expression in *S. woodyi* cells

*S. woodyi* cells were grown aerobically or anaerobically in the MR medium containing 20 mM DMSO and appropriate inductor for up to 4 h for quantitative reverse transcription polymerase chain reaction (RT-qPCR) assays or for up to 14 h for measurements of acrylate reductase activity.

RNA was extracted from *S. woodyi* cells using an RNA Solo kit (Evrogen) and additionally digested with RNase-free DNase I (Thermo Scientific, USA) at 37°C for 1 h. cDNA was synthesized using the MMLV RT kit (Evrogen) with random decanucleotide primers. A control reaction without reverse transcriptase was included for each sample. RT-qPCR assays were performed with a qPCRmix-HS SYBR kit (Evrogen), using the cDNA preparations as templates and A1/A2 and D3/D4 primer pairs (Table S1) for *ard*A and *ddd*Y, respectively. 16S rRNA was used for normalization (primer pair 16s_FWS/16s_RV). Serial dilutions of *S. woodyi* genomic DNA, which contains the genes for DddY, ArdA, and 16S rRNA in a 1:1:10 ratio, were used for calibration.

### Bioinformatics

Genomic context analysis of the *ddd*Y genes was performed using the webFlags program [16]. Multiple sequence alignment was done with Clustal Omega [17]. Cellular localization of bacterial proteins was predicted using SignalP 6.0 [18]. Operon identification was done with *Operon-mapper* [19], prediction of bacterial transcriptional terminators was performed using iTerm-PseKNC [20]. Protein identification by mass spectroscopy was carried out by MS+MS/MS ion search, using Mascot software version 2.3.02 (Matrix Science, USA) through the Home NCBI Protein Database.

## RESULTS

### 1. Isolation of acrylate reductase from periplasm of *S. woodyi*

For several reasons the luminescent marine bacterium *S. woodyi* [21] was considered to be a favorable model organism to identify acrylate reductase (Ard) of the anaerobic respiratory chain. *S. woodyi* genome contains the above-described *ddd*Y-associated gene cluster (Figs. 2a and S1), the bacterium is capable of DMSP cleavage and exhibits high acrylate reductase activity when grown anaerobically in the presence of acrylate (see Results, section 4).

When the periplasmic fraction from *S. woodyi* cells grown anaerobically in the presence of acrylate was separated by ion-exchange chromatography on DEAE Sepharose, a single peak of acrylate reductase activity was observed (Fig. 3a), which overlapped with one of the cytochrome peaks detected by light absorption at 405 nm.

**Figure 3.**
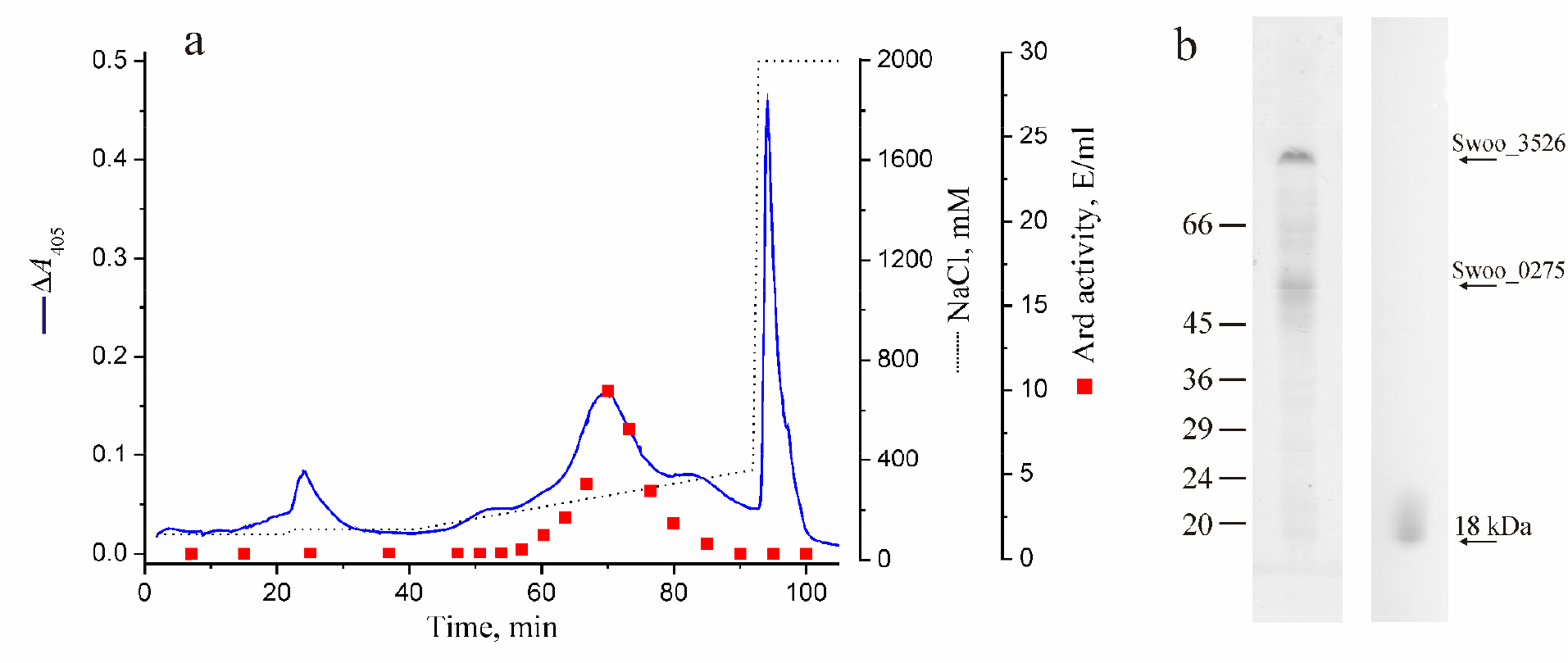
Ard isolation from *S. woodyi* cells. (a) Ion-exchange chromatography on DEAE Sepharose of a periplasmic fraction from *S. woodyi* cells grown anaerobically in the presence of acrylate. The blue line shows absorbance at 405 nm; the dotted line represents the NaCl concentration. Acrylate reductase activity in the eluate fractions is shown by red squares. (b) SDS-PAGE of the Ard preparation. The gel was stained for protein with Coomassie Blue (left lane) or for heme C with tetramethylbenzidine/H_2_O_2_ (right lane). Bars with numbers on the left side denote the positions and molecular masses of marker proteins. The protein bands identified by MALDI-MS analysis are marked on the right side.

SDS-PAGE analysis of the purified Ard preparation revealed two major protein bands with masses of ≈ 80 and 52 kDa (Fig. 3b, left panel). MS and MS/MS analyses of these bands indicated that the 80 kDa band was the trimethylamine N-oxide reductase TorA (Swoo_3526, sequence coverage: 50%), while the 52 kDa band was identified as the flavin-containing subunit of flavocytochrome *c* (Swoo_0275, sequence coverage: 65%). The latter finding confirms the prediction that Ard should include this protein as a flavin-containing subunit.

HPLC analysis of the Ard preparation for flavins (Fig. 4a) revealed only FAD (5 nmol mg^-1^). This finding provides support for the proposed Ard identification as all known flavocytochromes *c* contain FAD as a prosthetic group [2-4, 22, 23]. The spectra of Ard in the visible range (Fig. 4b) indicated the presence of cytochrome(s) *c*. Importantly, the contaminanting periplasmic protein TorA contains neither flavins nor hemes C [24]. Staining the electrophoregram of the Ard preparation for heme C reveals a heme-containing band with a mass of ≈ 18 kDa (Fig. 3b, right panel). However, MS and MS/MS analyzes of this band did not allow identification of the corresponding protein. A likely explanation is that the four CxxCH motifs of putative cytochrome subunit of Ard (Swoo_0276) may form eight covalent bonds with four hemes, limiting the number of unmodified tryptic peptides to one, located at the N-terminus of the mature protein. Given the uncertainty in the leader peptide cutting off place, the amino acid sequence of the N-terminal peptide could not be unambiguously predicted.

**Figure 4.**
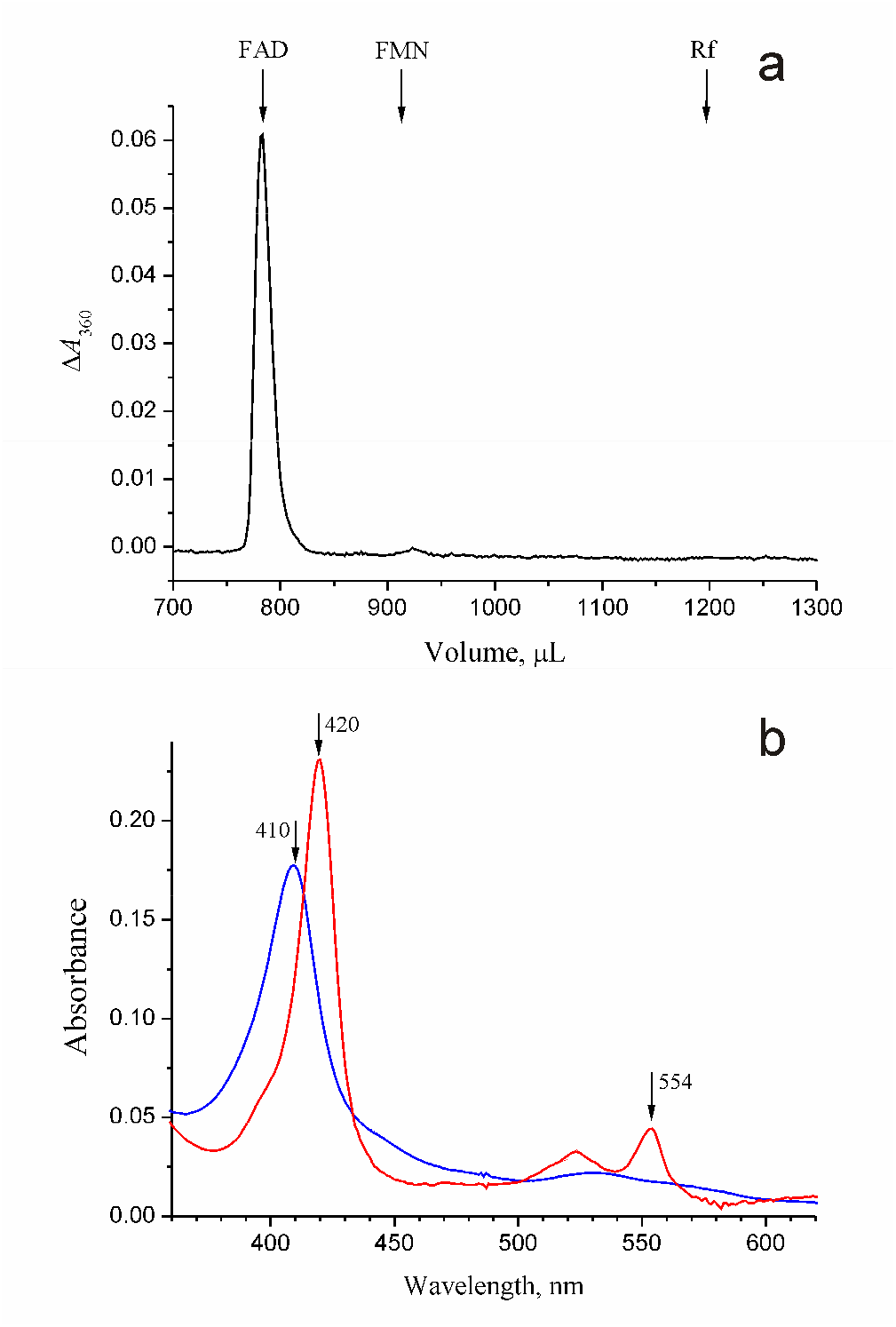
Identification of prosthetic groups in Ard. (a) HPLC separation of non-covalently bound flavins. The retention volumes for authentic FAD, FMN, and riboflavin (Rf) are indicated by arrows. (b) Electronic absorption spectrum of air-oxidized (blue curve) and dithionite-reduced (red curve) Ard. The spectra were measured in 100 mM Tris-HCl (pH 8.0) at 0.1 mg/mL Ard concentration. Cytochrome *c*-specific maxima of the γ- and α-bands of absorption are indicated by arrows.

### 2. Heterologous expression of the *S. woodyi swoo*_0275/*swoo*_0276 genes in *S. oneidensis* MR-1

The results of the above analysis of the periplasmic fraction of *S. woodyi* cells tentatively identified Ard of this bacterium as the Swoo_0275/Swoo_0276 flavocytochrome *c*. However, the presence of an additional, TorA protein in the isolated preparation and the inability to identify Swoo_0276 using mass spectrometry necessitated use of alternative approaches to support this identification. We therefore performed heterologous expression of the *swoo*_0275 gene or the pair *S. woodyi swoo*_0275/*swoo*_0276 genes in *S. oneidensis* MR-1 cells. The *S. oneidensis* MR-1 cells grown in the presence or absence of acrylate under either aerobic or anaerobic conditions demonstrated no periplasmic acrylate reductase activity (Table 1). This activity appeared only in *S. oneidensis* cells co-expressing a *swoo*_0275/*swoo*_0276-containing plasmid. Importantly, no acrylate reductase activity was detected in the *S. oneidensis* cells expressing a single gene of the flavin subunit (*swoo*_0275) (Table 1). These data provided thus strong support for the *S. woodyi* Ard identification as a flavocytochrome *c* consisting of the FAD-containing subunit Swoo_0275 (ArdA) and the heme C-containing subunit Swoo_0276 (ArdB).

**Table 1.** Ard specific activities in the periplasmic fraction of different *S. oneidensis* strains grown aerobically in the absence of acrylate.

| Strain | Acrylate reductase activity<sup>1</sup><br>(nmol min<sup>-1</sup> mg of cell protein<sup>-1</sup>) |
| --- | --- |
| *S. oneidensis* MR-1 | < 0.3 |
| *S. oneidensis* / pSwoo_0275&Swoo_0276 | 19 ± 4 |
| *S. oneidensis* / pSwoo_0275 | < 0.3 |

<sup>1</sup>From two independent experiments.

### 3. Substrate specificity of Ard

The reductase activity of Ard towards different natural α,β-unsaturated carboxylic acids was measured at their 1 mM concentration. Ard was found to be quite specific and capable of reducing only acrylate and methacrylate, but not crotonic, fumaric, sorbic, urocanic, cinnamic, *p*-coumaric, caffeic, or ferulic acids. Ard activity required the presence of a carboxyl group in substrate, as this enzyme did not reduce acrylamide. The Ard activity was maximal at pH ≈ 7.5.

The kinetic parameters of the Ard-catalyzed reduction of acrylate and methacrylate were determined. In both cases, a hyperbolic dependence of the rate of the catalyzed reaction on the substrate concentration was observed with *K*_m_ of 16 ± 0.4 μM for acrylate and 19 ± 1.1 μM for methacrylate. The corresponding maximal activity values were 58 ± 0.5 and 2.6 ± 0.1 μmol min^-1^ mg^-1^ (Fig. 5a). The similarity in the *K*_m_ values and the marked difference in the *V*_max_ values suggested that methacrylate may act as a competitive inhibitor of the acrylate reductase activity of Ard. Indeed, methacrylate suppressed the reductase activity of the enzyme in dose-dependent manner (Fig. 5b). The other α,β-unsaturated carboxylic acids tested (see above) did not inhibit the acrylate reductase activity of Ard.

**Figure 5.**
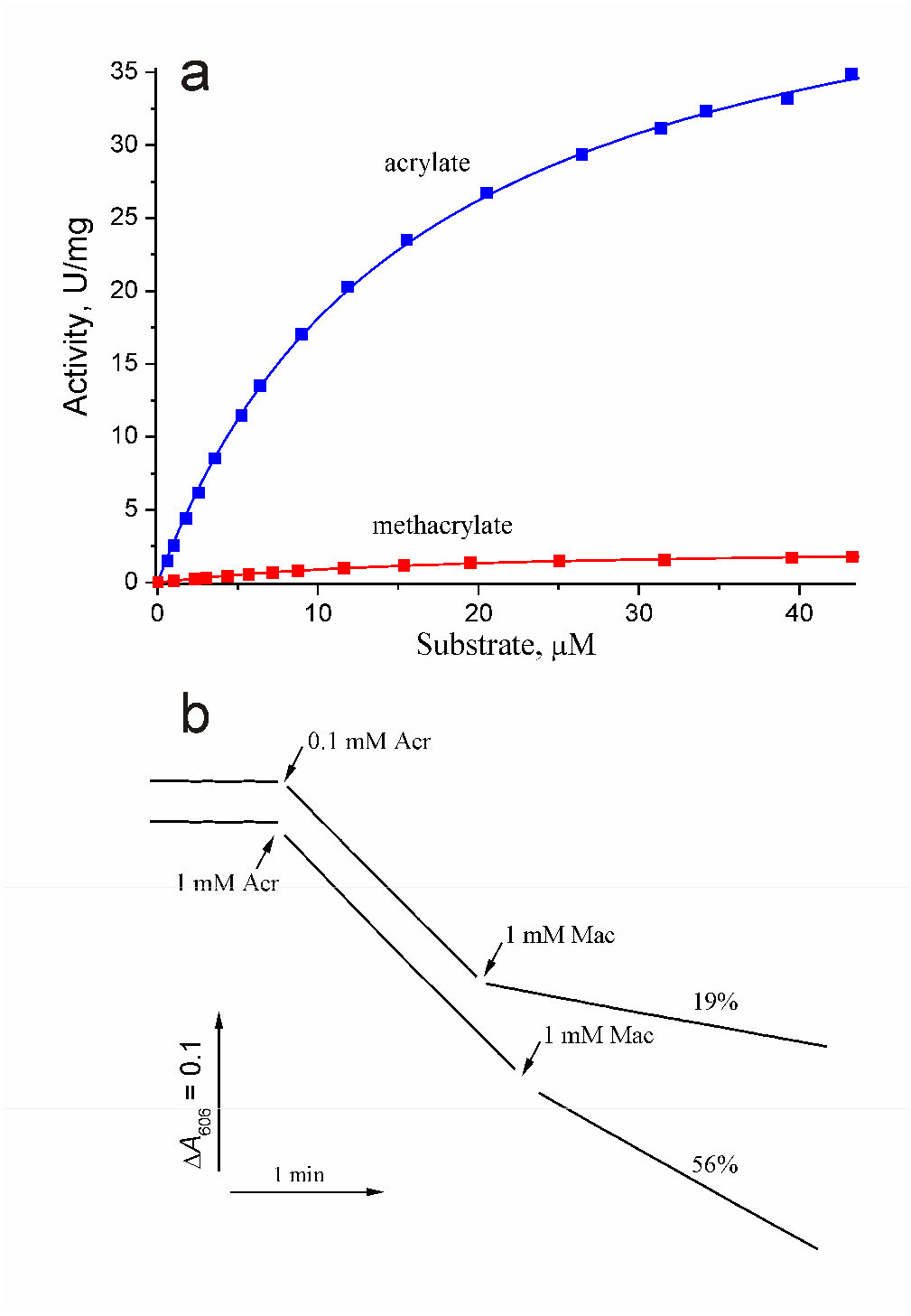
Enzymatic properties of Ard from *S. woodyi*. (a) Dependence of Ard activity on the concentration of acrylate (blue squares) or methacrylate (red squares). The lines show the best fits of the Michaelis-Menten equation. (b) Typical traces of MV oxidation by Ard. Arrows indicate additions of 1 or 0.1 mM acrylate (Acr) and 1 mM methacrylate (Mac). The numbers above the lines indicate enzymatic activity after methacrylate addition, where the activity before it is taken as 100%.

The ability of Ard to reverse acrylate reduction was also tested. This protein could not oxidize propionate with phenazine metasulfate and dichlorophenolindophenol as electron acceptors. Thus, Ard, like many other reductases of unsaturated carboxylic acids [1, 2, 4, 12, 23], catalyzes a unidirectional reaction and functions as a molecular diode [25].

### 4. Induction of Ard synthesis in *S. woodyi* cells

To determine the ability of the Ard substrates to induce its synthesis in *S. woodyi* cells, this bacterium was grown under aerobic or anaerobic conditions in the absence or presence of 1.5 mM acrylate or methacrylate or 5 mM DMSP. The selected (meth)acrylate concentration was the maximal acrylate concentration at which anaerobic growth was still observed. As Table 2 highlights, acrylate reductase activity was not detected in *S. woodyi* cells grown in aerobic conditions, even in the presence of the potential inducers. In contrast, a low but measurable acrylate reductase activity was detected in the cells grown anaerobically even in the absence of the inducers. The activity was further moderately increased in the presence of DMSP and significantly increased in the presence of acrylate or methacrylate (∼ 30- and 80-fold, respectively).

**Table 2.** Acrylate reductase activity and transcription of the *ard*A and *ddd*Y genes in *S. woodyi* cells grown under different conditions.

| Growth conditions | Acrylate reductase activity<br>(nmol·min<sup>-1</sup>·mg<sup>-1</sup>) | *ardA* mRNA/rRNA × 10<sup>-6</sup> | *dddY* mRNA/rRNA × 10<sup>-6</sup> |
| --- | --- | --- | --- |
| +O<sub>2</sub>, without inductors | <0.3 | 1.5 ± 0.2 | 3.2 ± 0.6 |
| +O<sub>2</sub>, 5 mM DMSP | <0.3 | 0.9 ± 0.1 | 2.7 ± 0.4 |
| +O<sub>2</sub>, 1.5 mM acrylate | <0.3 | 1.6 ± 0.3 | 3.5 ± 0.2 |
| +O<sub>2</sub>, 1.5 mM methacrylate | <0.3 | 0.9 ± 0.2 | 1.9 ± 0.1 |
| -O<sub>2</sub>, without inductors | 9.0 ± 4.0 | 20 ± 2.0 | 2.6 ± 0.2 |
| -O<sub>2</sub>, 5 mM DMSP | 45 ± 15 | 13 ± 1.0 | 1.7 ± 0.2 |
| -O<sub>2</sub>, 1.5 mM acrylate | 260 ± 30 | 87 ± 5.0 | 21 ± 1.0 |
| -O<sub>2</sub>, 1.5 mM methacrylate | 780 ± 50 | 220 ± 40 | 27 ± 2.0 |

As expected, *swoo*_0275–*swoo*_0277 gene cluster transcription pattern paralleled that of acrylate reductase activity. The programs *Operon-mapper* and iTerm-PseKNC predicted that the *swoo*_0275–*swoo*_0276 genes form an operon (*ard*), which does not include the *ddd*Y gene (*swoo*_0277). Therefore, transcriptional regulation was determined for both an *ard*-operon gene (*ard*A) and *ddd*Y. As Table 2 highlights, the *ard*-operon was not induced under aerobic conditions. In contrast, anaerobic growth led to a ∼13-fold increase in *ard* transcription and acrylate or methacrylate further stimulated it 4–11-fold.

Transcription of the *ddd*Y gene (*swoo*_0277) was induced in a similar manner. The maximal level of the *ddd*Y transcription was observed under anaerobic conditions in the presence of acrylate or methacrylate (Table 2). However, in the case of the *ddd*Y gene, the maximal effect was exerted by unsaturated carboxylic acids and anaerobiosis was, although a necessary, but not sufficient to increase the *ddd*Y transcription.

To determine the ability of *S. woodyi* to use Ard substrates as terminal electron acceptors for anaerobic respiration, cells were grown in the presence of acrylate, methacrylate, or DMSP, using the classical electron acceptor DMSO as a control. Acrylate in concentrations exceeding 1 mM was found to inhibit anaerobic growth, in line with known toxicity of this compound [10, 26]. Since terminal electron acceptor concentrations of less than 1 mM do not allow detection of growth stimulation during anaerobic respiration, further experiments were carried out using 10 mM methacrylate, which is much less toxic [27], and DMSP or DMSO at the same concentration. All three compounds increased growth yield by 20–25% (Fig. 6).

**Figure 6.**
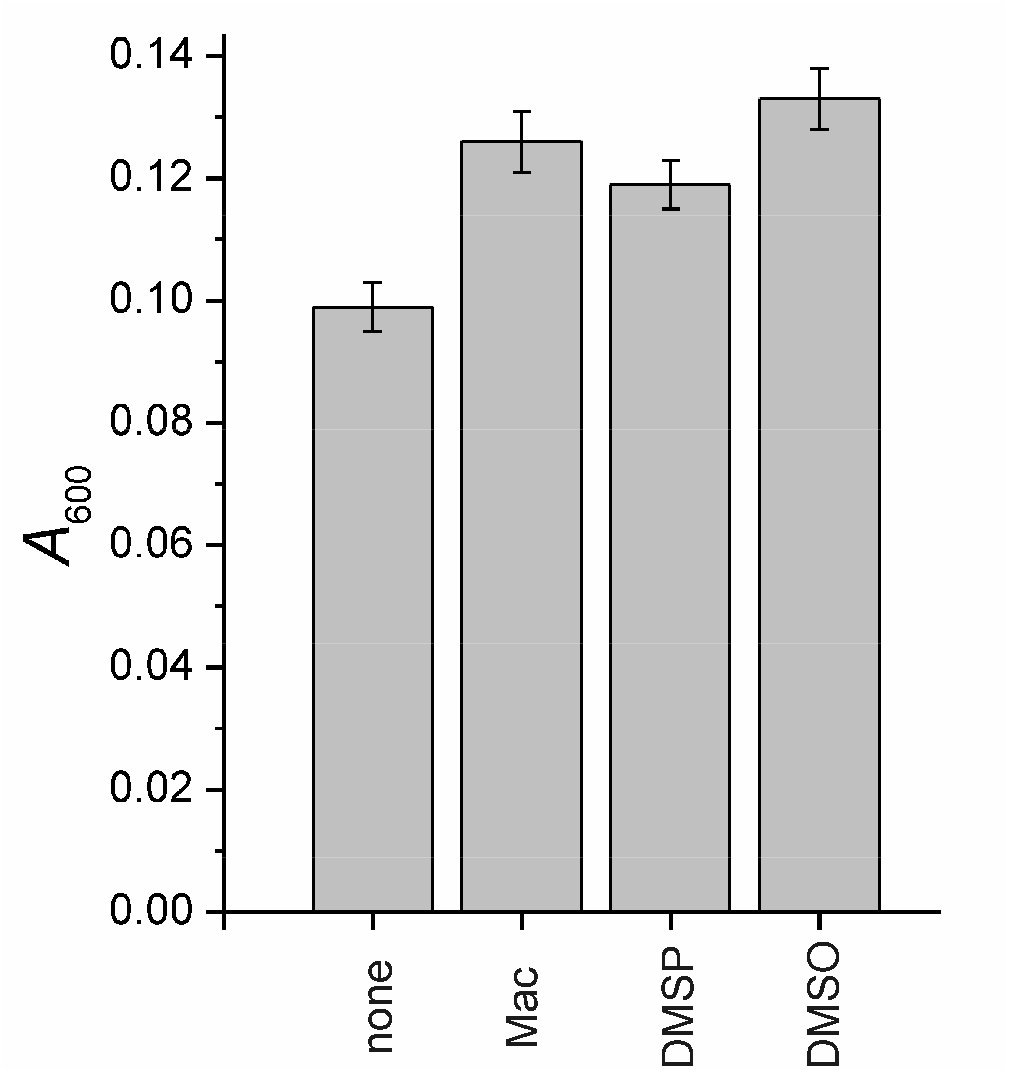
Anaerobic growth of *S. woodyi* in the presence of different electron acceptors. The initial light absorbance of the growth media at 600 nm was 0.005. The cells were grown anaerobically for 18 h at 25°C. Where indicated, the growth medium was supplemented with 10 mM methacrylate (Mac), DMSP, or DMSO. The bars show final absorbances with the error bars representing standard deviations in three independent experiments.

## DISCUSSION

The results reported above indicated that (i) acrylate reductase activity co-purifies with ArdA (Swoo_0275), the main protein of periplasmic fraction, when it is further fractionated; (ii) heterologous expression of the *ard*A/*ard*B genes of *S. woodyi* (*swoo*_0275/*swoo*_0276), but not of a single *ard*A gene (*swoo*_0275) in *S. oneidensis* MR-1 cells confers to them acrylate reductase activity of periplasmic localization, otherwise absent in authentic cells; (iii) expression of the *ard*-operon in *S. woodyi* is induced by acrylate and methacrylate under anaerobic conditions and is accompanied by the appearance of periplasmic acrylate reductase activity. Taken together, these observations indicate that the flavocytochrome *c* ArdAB (Swoo_0275/Swoo_0276) is responsible for acrylate reduction in *S. woodyi* cells.

Ard is a highly specific enzyme, demonstrating activity only against acrylate and methacrylate. Substrate screening has established the structural requirements for Ard substrates: they should be α,β-unsaturated carboxylic acids with small-size substituents in the α- and β-positions. Ard is more tolerable to the substituent size in the α-position than in the β-position, since Ard did not reduce crotonate (β-methylacrylate) but reduced methacrylate (α-methylacrylate), although at a lower rate compared with acrylate. While only slightly decreasing the strength of substrate binding to Ard, the methyl group in the α-position apparently mispositioned methacrylate in the active site, significantly slowing down hydride ion or proton transfer in methacrylate reduction by a factor of 22, compared with acrylate reduction (Fig. 5).

Ard synthesis in *S. woodyi* cells was maximal under anaerobic conditions in the presence of acrylate and, especially, methacrylate (Table 2). The greater inducing effect of methacrylate is probably non-physiological and could be explained by a slower depletion of this compound in the growth medium because of the much lower activity of Ard towards methacrylate. Low activity of DMSP as Ard inducer may similarly result from low steady-state concentration of acrylate, potent inducer, in the DMSP-containing growth medium because of greater Ard induction compared to DddY induction at anaerobic conditions. Noteworthy, high ArdAB/DddY ratio could help to avoid the toxic effect of acrylate during DMSP-dependent anaerobic growth of *S. woodyi*.

The acrylate-dependent induction of Ard synthesis under anaerobic conditions and its lack under aerobic conditions indicate that *S. woodyi* cells exploit this enzyme for anaerobic respiration with acrylate as a terminal acceptor for the electron transport chain. Experimental testing of this hypothesis is complicated by the inability of this bacterium to grow anaerobically in a minimal medium. In rich media, this bacterium could grow anaerobically even in the absence of electron acceptors, masking significantly their stimulation effect on growth. We failed to detect growth stimulation of *S. woodyi* under anaerobic conditions in the presence of acrylate (data not shown). As noted above, this could result from low concentration of acrylate (1 mM) used because of its toxicity [10, 26]. However, in the presence of 10 mM methacrylate or DMSP, a small but significant increase in growth yield (by 20–25%) was observed, comparable to the effect of a classical terminal acceptor DMSO (Fig. 6).

Genes of Ard-like proteins are widely distributed among various marine bacteria and form characteristic gene clusters with *ddd*Y on the chromosome (Fig. S1). The cluster allows anaerobic respiration on DMSP, a widespread component of marine habitat. The genome of *H. aestuarii*, the only known bacterium capable of using DMSP-originated acrylate as a terminal electron acceptor [7], also contains such a cluster, including the gene for an ArdA-like protein (GenBank: SHJ73509). The amino acid sequence of this protein is similar to that of *S. woodyi* Ard (50% identity, 66% similarity), with almost identical amino acid residues of their catalytic sites (Fig. 2b). Furthermore, the *ddd*Y-*ard*AB gene cluster (*so*_3622–*so*_3624) is also found in *S. oneidensis* MR-1 genome but is inactivated by the insertion sequence element ISSod3 [28]. This is apparently related to adaptation of this freshwater bacterium to life in conditions in which algae and plants do not synthesize DMSP because of low osmolarity of the environment.

## Conclusion

In the present work, the enzyme responsible for acrylate reduction in the electron transport chain of *S. woodyi* was identified. This protein enables DMSP-dependent anaerobic respiration in this and, possibly, other marine bacteria.

## Abbreviations

Ard: acrylate reductase
DMSO: dimethyl sulfoxide
DMSP: dimethylsulfoniopropionate
HPLC: high performance liquid chromatography
MS: mass spectrometry
MV: methyl viologen
RT-qPCR: quantitative reverse transcription polymerase chain reaction

## Acknowledgments

MALDI MS facility became available to us in the framework of the Moscow State University Development Program PNG 5.13.

