## Supplemented Figure S1 and Supplemented Table S1 for "Acrylate reductase of an anaerobic electron transport chain of the marine bacterium *Shewanella woodyi*"

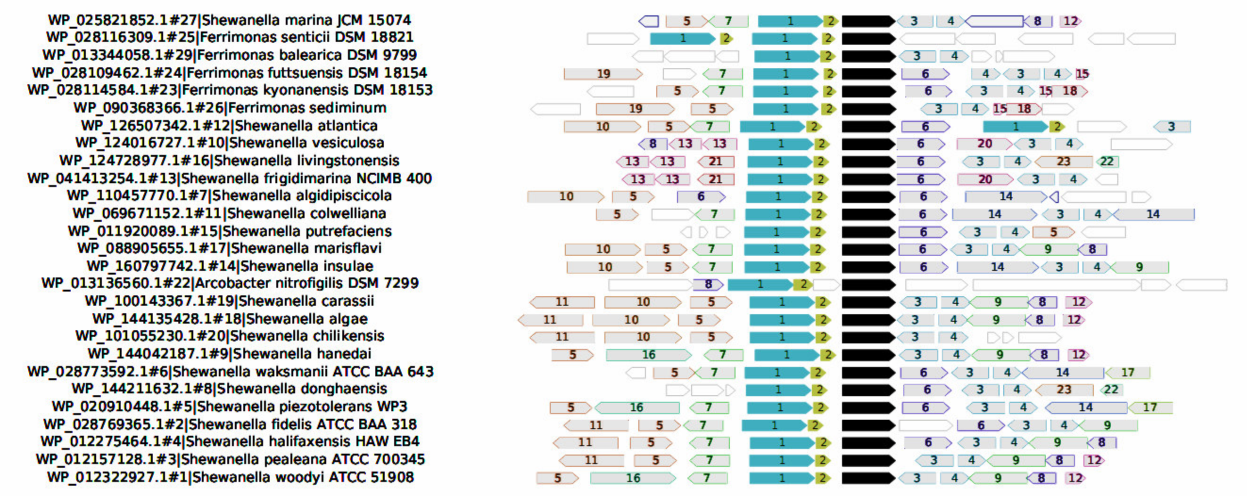


**Fig. S1**. **Genomic context analysis of the *ddd*Y gene (shown on the right panel by black boxes) in different *Shewanella* species and related bacteria.** The analysis was performed by the webFlaGs service [Saha *et al.*, 2021]. Bacterial species and accession numbers of corresponding DddY proteins are shown on the left panel. Genes encoding homologous proteins are marked by equal numbers inside boxes:

1 – flavoprotein subunit of flavocytochrome *c*;
2 – cytochrome $c_3$ family protein;
3 – LysR family transcriptional regulator;
4 – TetR/AcrR family transcriptional regulator;
5 – NADPH:acrylyl-CoA reductase;
6 – hypothetical protein;
7 – LysR family transcriptional regulator;
8 – response regulator;
9 – ATP-binding protein;
10 – $\sigma_{54}$-dependent Fis family transcriptional regulator;
11 – aldehyde dehydrogenase family protein;
12 – Spy/CpxP family protein refolding chaperone;
13 – hypothetical protein;
14 – methyl-accepting chemotaxis protein;
15 – DUF2282 domain-containing protein;
16 – hypothetical protein;
17 – inorganic phosphate transporter;
18 – DUF692 domain-containing protein;
19 – DUF885 domain-containing protein;
20 – anaerobic C4-dicarboxylate transporter;
21 – hypothetical protein;
22 – hypothetical protein;
23 –glutathione-disulfide reductase.

**Table S1.** Primers used in the work

| Primer | Sequence |
|---|---|
| Sh_wood_dir | 5’-CGCTTGGCTGCCTGAGTG-3’ |
| Sh_wood_CR4_rev | 5'-GGTTTACGTATCAATTTCTTACAA-3’ |
| A1 | 5'-AAGCAAGAGGGCGTTAAAGA-3’ |
| A2 | 5’-GTTTGGTAAGTACGTGCTACGG-3’ |
| D3 | 5'-ATGCCTGCGGTATTCCCACA-3’ |
| D4 | 5'-CTGCACCGACCTTTTCACCA-3’ |
| 16s_FWS | 5'-CAGCCACACTGGGACTGAGA-3’ |
| 16s_RV | 5’-GTTAGCCGGTGCTTCTTCTG-3’ |
